# Sex differences in choice-based thermal nociceptive tasks in adult rats

**DOI:** 10.1101/2021.11.28.470257

**Authors:** JR Bourgeois, AM Kopec

**Affiliations:** Dept. of Neuroscience & Experimental Therapeutics, Albany Medical College, Albany, NY

## Abstract

Interest in the role of sex as a biological variable continues to increase, including a mandate for the study of both sexes in NIH-funded research. Choice-based thermal nociceptive tests allow for the study of a more spontaneous response to thermal stimuli and avoidance behavior compared to traditional nociceptive assays, and their usage has been increasing in recent years. However, to date no comparison of naïve male and female responses to such tests has been published. As sex differences are known to exist in both human chronic pain conditions and rodent models of nociception, it is critical to understand the impact of sex on any nociceptive assay. Herein, we examined the effect of sex on two choice-based thermal nociceptive tests, the thermal gradient test and the temperature place preference test, in adult rats. We report that marked sex differences exist in responses to these tests. Namely, the activation of a 10° C-to-47° C thermal gradient results in an increase in time spent in the 10° C zone in females, compared to a reduction in males. In a temperature place preference test pairing a surface temperature of 22° C with either 5° C, 10° C, 47° C, or 50° C, males spent less than 50% of their time in every non-22° C zone, but in females this was only observed when testing 50° C. Together, these results suggest that male rats show more avoidance behavior to non-ambient temperatures when given free access to multiple zones, including at temperatures which are milder than those typically used to evoke a nociceptive response in traditional hot and cold plate tests.

## INTRODUCTION

The treatment of chronic pain remains a public health challenge, with approximately 20.4% of Americans experiencing chronic pain [1], defined as pain lasting longer than three months [2]. There are numerous tasks of thermal nociception with which to study animal models exhibiting a pain phenotype, with the hot plate test being among the most frequently used. The hot plate test is typically used to dissect the basic mechanisms of thermal nociception, to model hyperalgesia or allodynia observed in painful conditions, or to assess the efficacy of analgesics [3–7]. The cold plate test is used to model cold-based nociception, although its use is less frequent than the hot plate test, and the resulting behavior is often less pronounced [8–11]. While these tests measure responsive behavior such as paw withdrawal latency, paw licking, or jumping, these outcomes rely on experimenter interpretation, introducing some subjectivity. Such responsive behaviors, while informative, are considerably simplistic compared to the interaction of biological, psychological, and social aspects of human pain perception, particularly as it applies to chronic pain [12–16]. Additionally, the surface temperatures required to evoke an obvious response are often relatively extreme, e.g. ≥ 50 °C or ≤ 5°C for hot and cold plate tests, respectively, necessitating the use of trial cutoff times to prevent tissue damage [3, 5, 6, 8, 9, 17].

As an expansion of and complement to hot/cold plate tests, alternative assays to evaluate thermosensation which incorporate an element of choice have become more common in recent years, including the thermal gradient test, the thermal place preference test, and the Rotterdam Advanced Multiple Plate (RAMP) method [18–24]. These choice-based assays make use of flooring with a minimum of two discrete thermal zones and almost always include a non-noxious ambient zone, to which the animal has free access. The measured outcome of choice-based thermal assays is usually time spent in each zone, rather than nocifensive behaviors as identified by experimenters. Choice-based thermal assays provide an opportunity to observe behavior at less extreme temperatures which are not typically identified as aversive.

Studies of pain in humans indicate sex differences in the presentation of numerous chronic pain conditions. Specifically, complex regional pain syndrome (CRPS), fibromyalgia, rheumatoid arthritis, migraine headaches, lower back pain, and irritable bowel syndrome, among others, are all reported more frequently in females [25–32]. Sex differences exist in both the presentation and severity of pain, with females more commonly reporting neuropathic pain lasting more than three months and a higher percentage describing their chronic pain as “significant” than males [33, 34]. Sex differences in the experience of thermal pain also exist in healthy humans, with females showing increased sensitivity to both hot and cold temperatures [35–40]. In rodent models of thermal pain, traditional hot plate studies give conflicting results in sensitivity to aversive heat stimuli, with some studies showing no difference and others showing either males or females as being more sensitive to hot temperatures [5, 17, 41–44]. To our knowledge, no direct comparisons between male and female rats in response to the cold plate test have been published. Although choice-based thermal nociception assays are increasingly used in use in rodent pain models, typically only one sex is tested.

Here, we show evidence that sex differences are present in behavioral responses to two choice-based thermal assays: the thermal gradient test and the two-temperature preference test. In addition, we find that choice-based thermal assays may uncover evidence of avoidance behavior at less extreme temperatures than are typically observed in traditional hot and cold plate tests.

## METHODS

### Animal model

Adult male and female Sprague-Dawley rats were purchased to be breeding pairs (Harlan/Envigo) and were group-housed with ad libitum access to food and water. Colonies were maintained on a 12:12 light:dark cycle (lights on at 07:00) and cages were changed twice per week. Litters were culled to a maximum of 12 pups between postnatal day (P)2 and P5, and at P21 pups were weaned into same sex pair-housing. To reduce the impact of any litter-specific effects, no more than two animals per sex were used from any single litter for the same experiment. All animals were approximately 3 months old at the time of experiments. Experiments and animal care were approved by the Institutional Animal Care and Use Committees at Albany Medical College.

### Thermal gradient test

All tests were completed between the hours of 07:00-11:00 or 15:00-19:00 to minimize sleep cycle disruption. Prior to any behavioral testing, animals were handled on 3 occasions of ~5 minutes each in the behavioral testing suite, and animals were given at least 30 minutes to acclimate to the behavioral suite prior to testing. The thermal gradient chamber (Bioseb, BIO-GRADIENT) is a 10 cm x 120 cm enclosed rectangular chamber with opaque walls and a metal floor. An area of ~10 cm x 12 cm on each end is positioned atop a cold or hot plate, creating surface temperatures of 10°C and 47°C, respectively, at the ends of the chamber floor. The remaining ~96 cm of the chamber floor is not in direct contact with an underlying temperature plate. Although the surface area within ambient zone was not uniform (as a result of proximity to either the hot or cold plates), areas adjacent to the hot or cold zones but not directly over the temperature plates quickly regressed to ambient temperatures which would not be considered noxious (see **Supplemental Fig. 1** for temperature distribution), and thus time spent within this area was binned. Animals were given open access within the chamber, and movement was tracked via overhead camera. On day 1, animals were placed in the chamber for 30 minutes with the entire chamber at ambient temperature to determine initial zone preference. On day 2, temperature plates were turned on to create a temperature gradient, and the test was repeated. Day 2 results are expressed as the percentage change of time spent within a given zone, compared to average baseline preference for that group.

### Thermal place preference test

Animals were placed on an apparatus (Bioseb, BIO-T2CT) containing two independently controlled temperature plates. One plate was maintained at a surface temperature of 22° C, while a second plate was set to a surface temperature of either 5°C, 10° C, 47° C, or 50° C. Animals were allowed to move freely between the plates for 10 minutes, which was tracked via overhead camera, to determine the time spent in each temperature zone. Results for each experiment are reported as three separate outcomes: time spent in non-22° C zone, number of entrances into the non-22° C zone, and average time spent in non-22° C zone per entrance.

### Statistics

No single animal was used for all experiments, but most were used for 3-4 of 5 possible experiments. Each animal was subjected to only one test per day. Animals with outcomes greater than two standard deviations above or below the mean were excluded as outliers. For the thermal gradient test, change from baseline within any temperature zone following thermal gradient activation was the outcome used to determine outliers, and all thermal gradient data from such animals were excluded from all analyses, resulting in the exclusion of two animals. For the thermal place preference test, all data from any animal were excluded at a given temperature if any of the outcomes were more than two standard deviations from the mean, resulting in the exclusion of 2 5° C tests, 2 10° C tests, 2 47° C tests, and 3 50° C tests. Thermal gradient tests were analyzed using two-way ANOVAs (sex x temperature) with Tukey’s multiple comparisons post-hoc test. Between sex and within-sex comparisons of thermal place preference tests were analyzed using two-way ANOVAs (sex x temperature) with Tukey’s multiple comparisons post-hoc test. Additionally, to determine whether a temperature evoked avoidance behavior, for each temperature we compared the time spent in the non-22° C zone for each sex with results expected by chance using a one sample t-test. Statistical significance is defined as *p*<0.05.

## RESULTS

### Sex differences in temperature preference in the thermal gradient test

In an initial 30-minute test with the entire thermal gradient at ambient temperature, there were no significant differences between or within sex for any zone (two-way ANOVA, *n*= 8-10 animals/group) (**Fig. 1A**). The following day, animals underwent another 30-minute test with the hot and cold zones activated, and the percent change in time spent in each zone compared to group baseline averages was calculated. A main effect of temperature (F_(2,48)_=15.74, *p*<0.0001, n=8-10 animals/group) and an interaction between sex and temperature (F_(2,48)_=12.10, *p*<0.0001) were observed (**Table 1**). In response to the activation of the thermal gradient, males increased time spent in the ambient zone, and decreased time spent in both the hot (*p*=0.0028, Tukey’s multiple comparisons post hoc analysis) and cold zones (*p*=0.0047). In females, activation of the thermal gradient resulted in decreased time spent in the hot zone and increased time spent in both the cold (*p*<0.0001) and ambient zones (*p*=0.0111). A significant difference in response to thermal gradient activation was observed between males and females, with females increasing time spent in the cold zone compared to decreased time in the cold zone in males (*p*=0.0009) (**Fig. 1B**).

**Figure 1.**
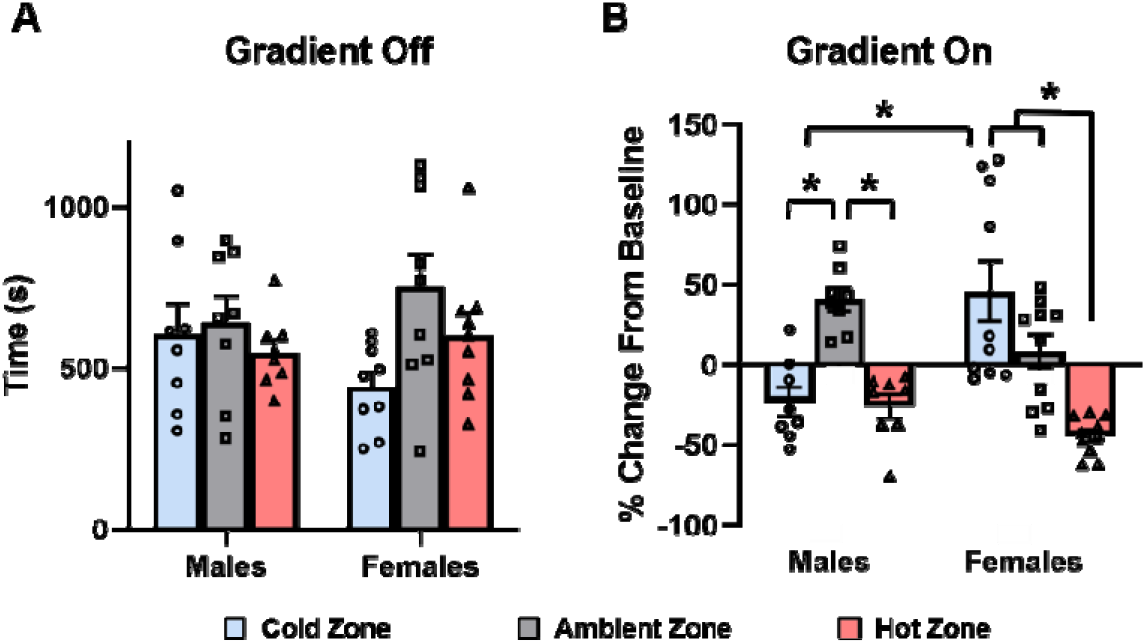
Sex differences in the thermal gradient test.

Males and females show different responses to the choice between hot, cold, and ambient thermal zones. (**A**) With the entire gradient at ambient temperature, no sex differences exist in the amount of time spent in each zone, nor is any within-sex preference observed between zones (two-way ANOVA, *n*=8-10 animals per group). (**B**) Following activation of the thermal gradient, males and females respond differently. Males reduced time spent in the cold and hot zones and increased time spent in the ambient zone, while females reduced time spent in the hot zone and increase time spent in the cold zone (*=*p*<0.05, two-way ANOVA with Tukey’s multiple comparisons post hoc analysis, main effect of temperature: *p*<0.001; interaction between sex and temperature: *p*<0.0001, *n*=8-10 per group).

**Table 1:** Statistical main effects and interactions.

| Assay | Outcome | Main Effect of Sex | Main Effect of Temperature | Interaction |
| --- | --- | --- | --- | --- |
| Thermal Gradient (Off) | Time in Zones | $F_{(1, 48)} < 0.0001$<br>$p = 0.9993$ | $F_{(2, 48)} = 2.195$<br>$p = 0.1224$ | $F_{(2, 48)} = 1.471$<br>$p = 0.2398$ |
| Thermal Gradient (On) | % Change in Preference | $F_{(1, 48)} = 0.4400$<br>$p = 0.5103$ | $F_{(2, 48)} = 12.10$<br>$p < 0.0001$ | $F_{(2, 48)} = 15.74$<br>$p < 0.0001$ |
| Thermal Place Preference | Time in Non-22°C Zones | $F_{(1, 71)} = 22.01$<br>$p < 0.0001$ | $F_{(3, 71)} = 15.63$<br>$p < 0.0001$ | $F_{(3, 71)} = 0.4921$<br>$p = 0.6889$ |
| Thermal Place Preference | Entrances into Non-22°C Zones | $F_{(1, 71)} = 12.86$<br>$p = 0.0006$ | $F_{(3, 71)} = 3.591$<br>$p = 0.0177$ | $F_{(3, 71)} = 0.2805$<br>$p = 0.8393$ |
| Thermal Place Preference | Time Per Entrance into Non-22°C Zones | $F_{(1, 71)} = 8.173$<br>$p = 0.0056$ | $F_{(3, 71)} = 11.30$<br>$p < 0.0001$ | $F_{(3, 71)} = 0.3027$<br>$p = 0.8234$ |

### Sex differences in temperature preference in the thermal place preference test

To test preferences when given a direct choice between two thermal zones, we then conducted a series of thermal place preference tests allowing animals free access for 10 minutes between a 22° C zone and relatively cold or hot zone. Animals were tested at surface temperatures of 10° C and 47° C, as were used in the thermal gradient test, as well as more extreme cold and hot temperatures (5° C and 50° C, respectively). Main effects of both sex (F_(1,71)_=22.01, *p*<0.0001, two-way ANOVA, n=9-10 animals/group) and temperature (F_(3,71)_=15.63, *p*<0.0001) were observed in the time spent in each non-22° C zone (**Table 1**). Males spent significantly less time in the 50° C zone compared to the 10° C zone (*p*=0.0008, Tukey’s multiple comparisons post hoc analysis). Comparing across all tests, males spent significantly less time in the 50° C zone compared to the 10° C zone (*p*=0.0008, Tukey’s multiple comparisons post hoc analysis), while females spent less time in the 50° C zone compared to the 5° C zone (*p*=0.0085), the 10° C zone (*p*<0.0001), and the 47° C zone (*p*=0.0168). Additionally, males spent less time than females in the 47° C zone (*p*=0.0419) (**Fig 2A**). Main effects of both sex (F_(1,71)_=12.86, *p*=0.0006) and temperature (F_(3,71)_=3.591, *p*=0.0177) were also observed in the number of entrances made into the non-22° C zones (**Table 1**), although no within-sex or within-temperature differences reached statistical significance after post-hoc analysis (**Fig. 2B**). Finally, main effects of sex (F_(1,71)_=8.173, *p*=0.0056) and temperature (F_(3,71)_=11.30, *p*<0.0001) were observed in the amount of time spent in each non-22° C zone per entrance (**Table 1**). Males showed a reduction in time spent per entrance in the 50° C zone compared to the 10° C zone (*p*=0.0350). Females showed a reduction in time spent per entrance into the 50° C zone compared to the 5° C zone (*p*=0.0076) and the 10° C zone (*p*=0.0012), but not the 47° C zone, although a non-significant trend was present in this comparison (*p*=0.0527) No sex differences were observed in time spent in thermal zones per entrance (**Fig. 2C**).

**Figure 2:**
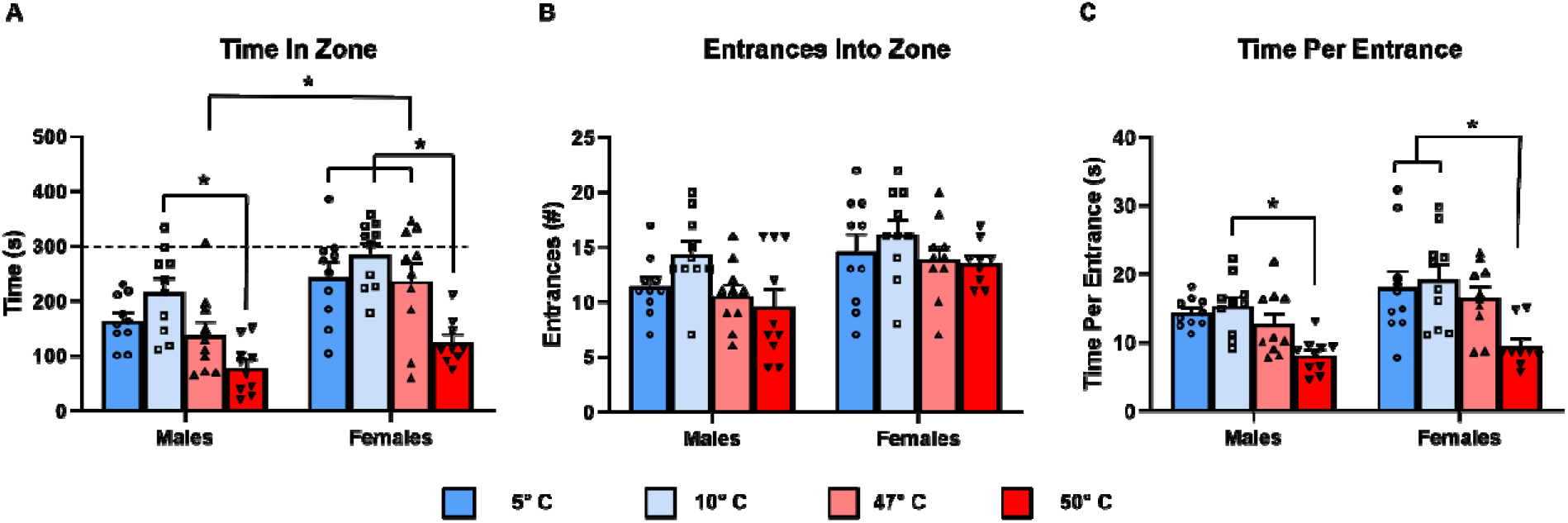
Sex differences in thermal place preference test at more extreme temperatures. In a 10-minute thermal place preference test, (**A**) males spent more time in the 10° C zone than the 50° C zone, while females spent more time in the 5° C, 10° C, and 47° C zones than the 50° C zone. Additionally, males spent less time in the 47°C zone than females. (**B**) No differences were observed in the number of entrances into any zone between males and females, nor between different temperatures in within-sex comparisons. (**C**) Males spent more time per entrance in the 10° C zone than the 50° C zone, while females spent more time per entrance in the 5° C and 10° C zones than the 50° C zone. (*=*p*<0.05, two-way ANOVA with Tukey’s multiple comparisons post hoc analysis, *n*=9-10 animals per group).

Between a 22° C zone and a 10° C zone, males spent less time in the 10° C zone than expected by chance (*p*=0.0080, one sample t-test, *n*=10 animals/group), while female results did not differ from chance (**Fig. 3A**). Similarly, when given the choice between a 22° C zone and a 47° C zone, males spent less time in the 47° C zone than predicted by chance (*p*<0.0001), while females did not (**Fig. 3B**). When given a choice between a 22° C and a 5° C zone, males spent less time in the 5° C zone than expected by chance (*p*<0.0001), while females did not (**Fig. 3C**). Finally, between a 22° C and a 50° C zone, both males and females spent less time in the 50° C zone than expected by chance (*p*<0.0001 for both, one sample t-test) (**Fig. 3D**).

**Figure 3:**
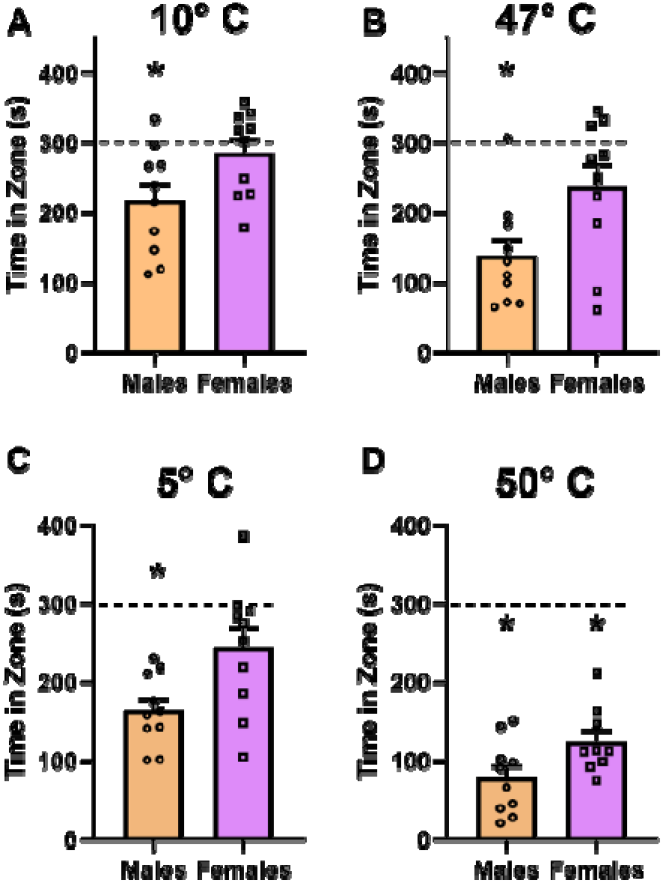
Sex differences in aversion in the thermal place preference test. In a 10-minute thermal place preference test, males spent less than 50% of their time in the (**A**) 10° C, (**B**) 47° C, and (**C**) 5° C zones when given the choice between these temperatures and a 22° C zone. Females did not show this response. (**D**) Both males and females spent less time in the 50° C zone than expected by chance (*=*p*<0.05, one sample t-test; *n*=9-10 animals per group).

## DISCUSSION

Herein, we examined the behavior of male and female adult rats in two different choice-based thermal tests. Consistent with their behavior in the thermal gradient test, males exhibited avoidance behavior of 10° and 47° C in the thermal place preference test, as well as more extreme 5° and 50° C. These results in male rats indicate aversive behavior at temperatures which are milder than those typically used in traditional hot and cold plate tests [3, 8–10, 45]. In contrast, females exhibited a cold preference in the thermal gradient test, and only avoided the 50° C zone in the thermal gradient test, indicating an overall greater tolerance for both hot and cold temperatures. Collectively, these data are particularly interesting when compared with thermal nociception data in humans, which indicate higher sensitivity in healthy females to both hot and cold temperatures [35–40].

While traditional hot and cold plate tests have proved useful over decades of research, such tests rely on observer interpretation of reaction to a thermal stimulus. In the hot plate test, the measured reaction is frequently latency to respond, which is typically identified by paw lifting/licking or jumping. Cold plate tests generally produce a less pronounced response, and as a result outcomes are frequently measured as total response behaviors over a set amount of time, often several minutes. Observer interpretation requires a clear all-or-none response, temperatures are often set at a level designed to produce this effect. As a result, the risk of tissue damage is present, particularly in the case of the hot plate test, which may use temperatures as high as 58° C. Cutoff times must be implemented to reduce this possibility, especially when testing analgesics which may prevent the response required to end the test [3, 5, 8, 11]. Additionally, the introduction of an unescapable (and usually extreme) nociceptive thermal stimulus introduces an element of stress for the animal, which may present as a confounding variable in interpreting the results [46, 47]. The use of choice-based thermal nociception assays can serve to supplement the vast existing body of research on thermal nociception by examining the effects of a thermal stimulus on typical exploratory behavior. Such assays may therefore be particularly relevant to the study of chronic pain in humans, which involves a more complex interaction of biological and behavioral factors. Indeed, given the known sex differences in pain perception in humans, animal models of human pain must consider that psychological and social factors may exacerbate any underlying sex differences in human pain biology [12–14]. For example, catastrophizing, a negative mental state brought on by both actual and anticipated pain which often accompanies human chronic pain states and can impact the experience of clinical and experimental pain, is reported more commonly in women than men [15, 16, 48]. In light of this, sex differences in choice-based thermosensation assays may become particularly relevant due to their focus on anticipation and avoidance of discomfort, rather than the supraspinal reflex and nocifensive responses typically studied with hot and cold plate tests [6, 49]. While a paw withdrawal or lick as a reaction to an unavoidable noxious stimulus can be used to indicate a nociceptive response, in isolation it provides no information as to how the presence of such a stimulus may influence future behavior. Thus, the operant response of stimulus avoidance can provide important insight into how males and females are affected differently in chronic pain models [50]. Therefore, discrepancies in results between nociceptive assays which rely on reflexive responses versus assays which rely on more proactive avoidance behavior, particular in the context of sex differences, may provide valuable insights and hypotheses into the mechanisms driving them. Our data provide the first comparison of naïve male and female rodent behavior in these tests.

### Sex differences in temperature-induced behavior

The thermal place preference test involves only two thermal plates which are placed directly adjacent, with the entire apparatus enclosed in a clear plexiglass box, allowing for an interpretation of how a single temperature can influence behavior when paired with a non-noxious temperature choice in rats. Using this assay at the relatively mild cold and hot temperatures of the 10° C and 47° C, respectively, clear sex differences were observed in the response. Specifically, females spent ~72.0% more time in the 47° C zone compared to males **(Fig. 2A)**. While males spent less than 50% of their time in both the 10° C and 47° C zone, in females the time spent in each of these zones was not significantly different from chance **(Fig. 3A, 3B)**. This suggests that these temperatures are considered aversive to males, but not females, when presented in the context of a choice.

Using the thermal place preference test to compare males and females at more extreme temperatures, such as those typically used in traditional cold and hot plate tests, sex differences were still observed. Similar to the results seen at 10° C, males spent less time in the 5° C zone than expected by chance, whereas females did not (**Fig. 3C**). Interestingly, this aversion shown by males to both 5° C and 10° C in the thermal place preference test is at odds with published literature on naïve adult male behavior in the cold plate test. Specifically, multiple studies indicate a lack of response by naïve adult males to a 10° C cold plate, considering it non-noxious, with 5° C [9, 51] or 3° C [8] identified as the cutoff for noxious cold. Similarly, Kato, et al. identified a 10° C cold plate test as appropriate for identifying cold allodynia, with 4° C appropriate for identifying hyperalgesia, in an oxaliplatin-induced cold hypersensitivity model [10]. However, when assessing naive adult rats in a thermal place preference test against a 25° C plate, Balayssac, et al. found 17° C as the cutoff below which rats would spend less time in the cold zone, with time spent in 5° C and 10° C zones being roughly equivalent to one another (but both below the time spent in 17° C) [52]. While Kato, et al. found evidence of oxaliplatin-induced allodynia (at 10° C) and hyperalgesia (at 4° C) using a cold plate test, Balayssac, et al. found no effect of oxaliplatin at either 5° C or 10° C using a thermal place preference test. This suggests that while an oxaliplatin treatment is sufficient to evoke a behavioral response (in Kato et al.’s case, hind paw withdrawal) in a cold plate test, it is not sufficient to alter behavior in a more spontaneous choice-based test. To the best of our knowledge, no data have been published on the cold temperatures considered noxious to naïve adult female rats.

In contrast to our results at 5° C, our thermal place preference test examining more extreme heat at 50° C shows aversive behavior in both sexes. Indeed, 50° C was the only temperature of four tested in which females spent less time in the non-22° C zone than expected by chance (**Fig. 3D**). This temperature also showed sex differences in terms of within-sex comparisons to other temperatures tested. Namely, when given the choice between a 22° C and a second zone with a temperature of either 5° C, 10° C, 47° C, or 50° C, males showed reduced time spent in the 50° C compared only to the 10° C zone, while females spent less time in the 50° C zone than in the 5°, 10°, and 47° C zones (**Fig. 2A**). Similarly, while the time spent in the 50° C zone per entrance was only reduced compared to 10° C, females showed reduced time spent per entrance in the 50° C zone relative to both 5° and 10° C (**Figure 3C**). The fact that females spent less than half of their time in the 50° C, while this was not the case with the 47° C zone, is unsurprising. Indeed, temperatures of ≥50° C are typically used for hot plate tests, with the more rarely used 48° C being considered the lower end of the range of appropriate temperatures [3]. However, this still leaves an interesting discrepancy between male and female responses to the thermal place preference test at both 47° C and 50° C, as male behavior in this assay indicated avoidance behavior at 47° C (below the temperatures typically used to induce a nocifensive response), whereas no such behavior was seen in females.

### Sex differences in response to competing interests

The thermal gradient apparatus lends itself to the study of competing aversive interests. The thermal gradient test allows animals open exploration of a metal floor, enclosed in an opaque chamber with a clear plexiglass lid. While this setup allows for animals to spend time in preferred temperature zones, the presence of enclosed corners at each end (corresponding with the 10° C to 47° C zones) adds the variable of differing levels of potential exposure. In the absence of varied thermal stimuli, it should be expected that rodents would choose to spend more time in regions that provide more cover, as is typically seen in open field and elevated plus-maze tests [53–55]. Indeed, when the thermal gradient is not activated, both males and females spend a disproportionately reduced amount of time in the area designated as the ambient zone. Specifically, males and females spent only 35.7% and 41.8% of their exploratory time, respectively, in the ambient zone, despite the surface area of this zone accounting for 80% of the gradient floor **(Fig. 1A; Supplemental Fig. 1).** As a result, when the gradient is activated, the thermal gradient test should not be thought of as a strictly thermal test, but rather one of competing preferences between non-aversive temperatures and areas that may be perceived to provide increased protection. Although no differences in time spent within a given zone were observed between sexes with the gradient off, the activation of a thermal gradient led to different responses based on sex. Specifically, males spent reduced time in both the cold and hot zones, increasing their time spent in the more exposed ambient zone. In contrast, females responded to thermal gradient activation by reducing their time spent in the hot zone while increasing their time spent in the cold zone **(Fig. 1B)**. This may be interpreted in different ways. First, it is possible that females find the level of exposure present in the ambient zone more aversive than males, causing them to tolerate the aversive cold zone. However, naïve adult females have previously been shown to be more willing than males to enter and remain in the center zone of in a large open field test, and in the open arms of an elevated plus maze test, as well as showing a reduction in overall fear and anxiety-like behavior, suggesting that their lack of a shift to the ambient zone may not be due to an increased fear of exposure [53–55]. Alternatively, it is possible that males find the 10° C cold zone more aversive than females do, and thus were more willing to increase their time in the more exposed ambient zone. This interpretation is supported by the differences in behavior observed between sexes in the 10° C vs. 22° C thermal place preference test. The fact that females spend equal time in both zones, together with their shift to the 10° C zone in an activated thermal gradient, indicates that the 10° C surface temperature in the thermal gradient is less aversive to females than increasing time in the more exposed ambient zone. The observed behavior of males shifting their preference to the more exposed ambient zone rather than the cold zone in the activated thermal gradient is also consistent with their behavior in the 10° C vs. 22° thermal place preference test, which collectively suggest the 10° C surface temperature is more aversive in males.

To summarize, we identified sex differences in responses to hot and cold temperatures in two choice-based thermosensation assays in naïve adult rats. These findings may aid in the design of future studies using choice-based tests to study nociceptive responses in various rodent pain models, as well as identifying how incorporating an element of choice in nociceptive assays may reveal additional differences in behavior beyond the traditionally studied withdrawal responses.

## Supplemental Figure 1

**Supplemental Figure 1:**
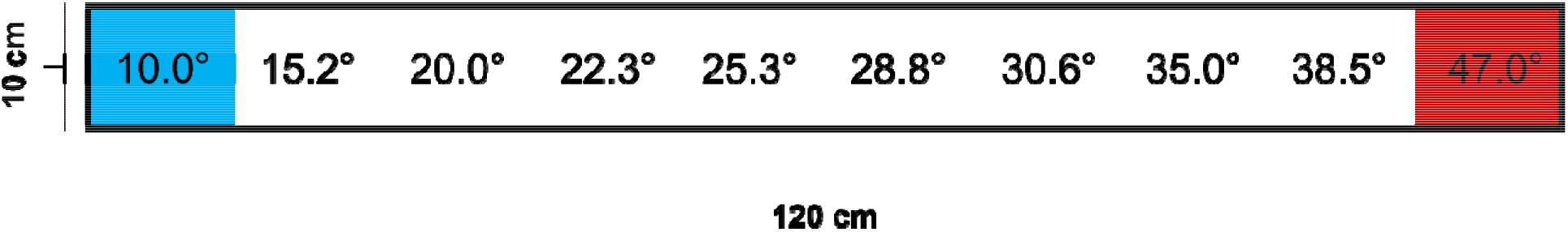
Schematic of thermal gradient chamber (top view) Representation of floor surface temperatures throughout the thermal gradient chamber, as depicted from above. Blue and red areas represent the portion directly overlaying cold and hot plates, respectively. The ambient zone (shown in white) was divided into 8 sections measuring 10 cm x 12 cm each, and surface temperature was obtained in the center of each section.

